# Microglia are involved in regulating histamine dependent and non-dependent itch transmissions with distinguished signal pathways

**DOI:** 10.1101/2022.09.23.498040

**Authors:** Yuxiu Yang, Bin Mou, Qi-Ruo Zhang, Hong-Xue Zhao, Jian-Yun Zhang, Xiao Yun, Ming-Tao Xiong, Ying Liu, Yong U. Liu, Haili Pan, Chao-Lin Ma, Bao-Ming Li, Jiyun Peng

## Abstract

Although itch and pain have many similarities, they are completely different in perceptual experience and behavioral response. In recent years, we have a deep understanding of the neural pathways of itch sensation transmission. However, there are few reports on the role of non-neuronal cells in itch. Microglia are known to play a key role in chronic neuropathic pain and acute inflammatory pain. It is still unknown whether microglia are also involved in regulating the transmission of itch sensation. In the present study, we used several kinds of transgenic mice to specifically deplete CX3CR1+ central microglia and peripheral macrophages together (whole depletion), or selectively deplete central microglia alone (central depletion). We observed that the acute itch responses to histamine, compound 48/80 and chloroquine were all significantly reduced in mice with either whole or central depletion. Spinal c-fos mRNA assay and further studies revealed that histamine and compound 48/80, but not chloroquine elicited primary itch signal transmission from DRG to spinal Npr1- and somatostatin-positive neurons relied on microglial CX3CL1-CX3CR1 pathway. Our results suggested that central microglia were involved in multiple types of acute chemical itch transmission, while the underlying mechanisms for histamine dependent and non-dependent itch transmission were different that the former required the CX3CL1-CX3CR1 signal pathway.

## Introduction

Pruritoceptive itch is an unpleasant sensation that is initiated from the skin and elicits a desire to scratch (Yosipovitch et al., 2003). Many kinds of stimuli upon or beneath skin can evoke itch sensation, such as a slight mechanical stroke, particular types of chemicals and inflammation induced cytokines. Histamine (HA) is one of the most well-known itch mediators that act on H1 and H4 receptors at the free-ending of pruriceptive nerve fibers (Dong and Dong, 2018; Kashiba et al., 1999; Strakhova et al., 2009). There are also several kinds of non-histamine itch mediators that used on laboratory animal studies, such as chloroquine (CQ), serotonin (5-HT), β-alanine and so on (Dong and Dong, 2018). Similar as pain sensation, chemical itch signals are known to be conveyed by unmyelinated C fibers (Ringkamp et al., 2011; Schmelz et al., 1997; Shim and Oh, 2008). Different groups of Dorsal Root Ganglion (DRG) neurons respond to those stimuli. Using single-cell RNA sequencing data, the DRG neurons could be categorized to 11 subpopulations. Among them, three non-peptidergic (NP) groups were associated with itch: NP1 expressing MrgprD responds to β-alanine, NP2 expressing MrgprA3 responds to several kinds of mediators including CQ, HA and BAM8-22, and NP3 expressing brain natriuretic peptide (BNP) and somatostatin (SST) responds to HA (Usoskin et al., 2015). Although itch and pain share some common peripheral afferent pathways and influence each other (Davidson and Giesler, 2010; Simone et al., 2004), the spinal level transmission pathways are quite different. Studies have revealed that itch transmission relies more on neuropeptides release rather than glutamate release from the peripheral neurons (Liu et al., 2010). Neuropeptides that were found to anticipate in itch signal transmission includes Gastrin-Releasing Peptide (GRP), Natriuretic Polypeptide B (NPPB), Neuromedin B (NMB), SST, substance P, et. al. (Akiyama et al., 2014; Huang et al., 2018; Mishra and Hoon, 2013; Wan et al., 2017).

Peripheral immune cells play important roles on chemical pruritogen insult detection and chronic itch development (Pasparakis et al., 2014). Mast cells are the major endogenous HA source (Rao and Brown, 2008). Type 2 T helper (Th2) cells, macrophages or dendritic cells and infiltrated monocytes are involved in cytokine release, including IL-4, IL-13 and IL-31 (Brandt and Sivaprasad, 2011; Nattkemper et al., 2018; Oetjen et al., 2017). Despite the well-known involvement of immune system in the periphery, whether immune cells participate in the central transmission of itch sensation is largely unknown.

Microglia are the resident immune cells in the central nerve system (CNS). Microglia share many properties with peripheral macrophages (e.g., expression of CX3CR1), but are originated from different source during early development (Gomez Perdiguero et al., 2015). A bunch of studies have revealed that microglia are one of the key players in pain hypersensitivity and chronic pain development (Inoue and Tsuda, 2018; Peng et al., 2016). With peripheral nerve injury, spinal microglia would be activated by multiple signals, including the fractalkine or CX3CL1 signals, through CX3CR1 receptors, ATP/ADP signals through purinergic receptors such as P2X4, P2X7 and P2Y12, and the CSF1 signals. These signal pathways are thought to be involved in microglial promoting pain hypersensitivities. To study whether microglia and peripheral resident macrophages are involved in acute itch sensation, and what is the signal pathway, we used multiple transgenic tools to deplete microglia and peripheral macrophages together or deplete central microglia alone, and examined how itch responses that induced by either HA dependent or non-dependent insults were affected. We demonstrated that central microglia were involved in both HA dependent and non-dependent (CQ) itch transmission. Further Spinal c-fos mRNA assay and behavior screening studies with the P2Y12 KO and CX3CR1 KO mice revealed that the underlying mechanisms for HA dependent and non-dependent itch transmission are different that the former requires the CX3CL1-CX3CR1 signal pathway, while the latter does not. The microglial P2Y12 receptors are not involved in any acute itch signal transmission.

## Results

### Depletion of CX3CR1+ microglia and macrophages inhibited acute itch responses to HA, C48/80 and CQ

The CX3CR1 fractalkine receptors are highly expressed in central microglia and peripheral macrophages. To establish an overall effect of CX3CR1+ cell depletion on acute itch transmission, we first used the CSF1R^f/f^;CX3CR1^CreER/+^ transgenic mice to knock out the csf1r gene in all CX3CR1+ cells. Because the CSF1 receptors in microglia and macrophages are essential for the cell survival (Elmore et al., 2014; MacDonald et al., 2010), knock out of the gene will cause cell apoptosis. With 3 doses of Tamoxifen (TM, 150 mg/kg, i.p., 48 hr interval) treatment, 98.4±1.1% of microglia in the spinal cord and 80.6 ± 2.9% of macrophages in the DRG were depleted when checked at 24 hr after the last Tamoxifen injection. While in the skin of the back neck, the number of F4/80+ labeled dendritic cell was not affected (Fig. 1A and B). Thus, the CSF1R strategy efficiently depleted the central microglia and the DRG macrophages, but did not affect peripheral dendritic cells.

**Figure 1.**
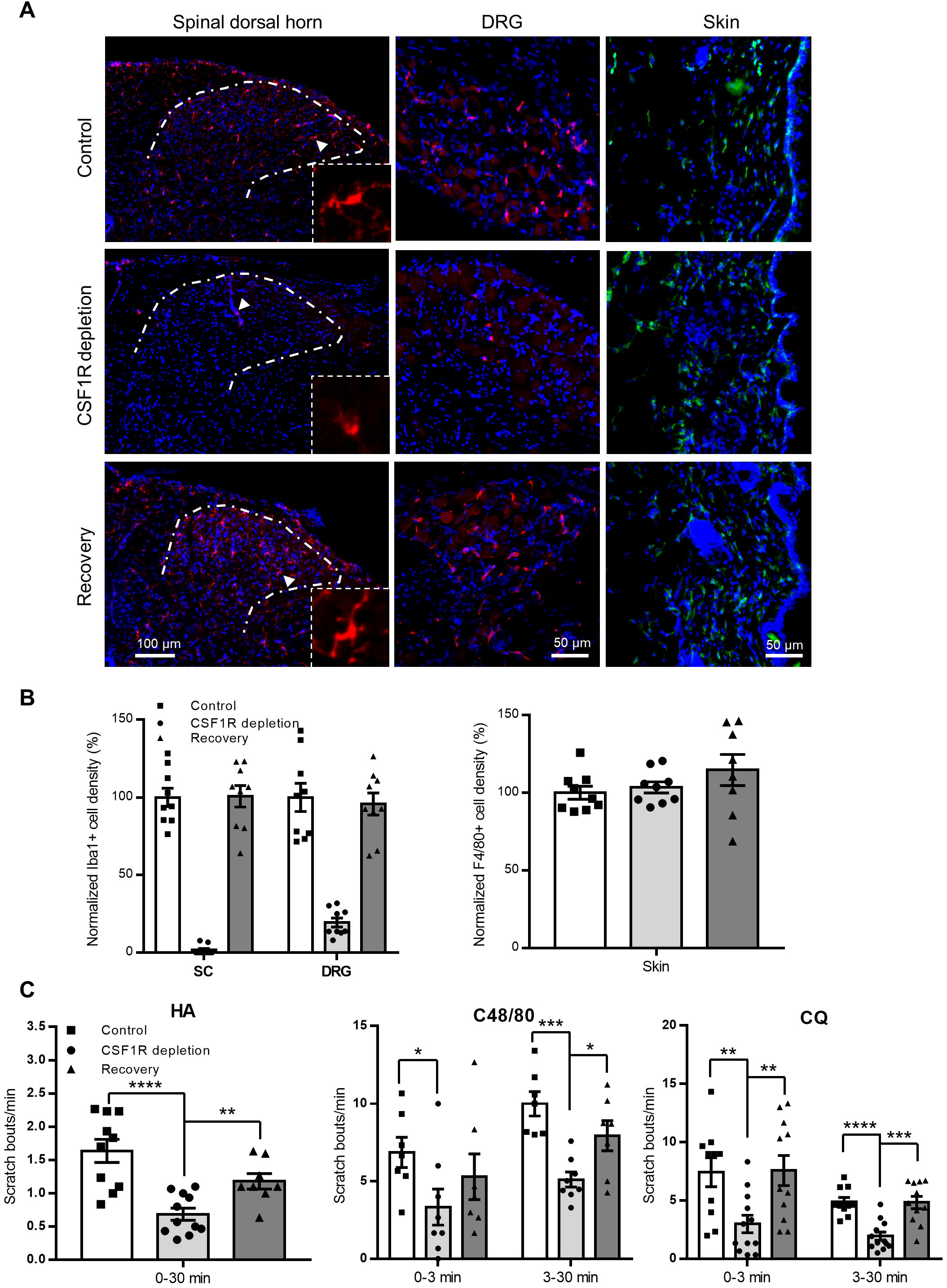
CSF1R depletion inhibited acute itch responses to HA, C48/80 and CQ. (**A-B**) With Tamoxifen (TM) treatment, the iba1+ microglia cells in spinal cord (SC) and macrophage cells in dorsal root ganglion (DRG) of the CSF1R^f/f^;CX3CR1^CreER/+^ mice were reduced to 1.6±1.1% and 19.4±2.9% of basal levels, respectively. While the F4/80+ dendritic cells in the skin were not altered. The recovery samples were obtained at 15 days after the last TM injection. Iba1+ cells in both SC (p = 0.938, un-paired t-test) and DRG (p = 0.721) were returned to control levels (n = 3 mice for each group, CSF1R^f/f^ mice were used as control and received the same doses of TM). The arrow indicated cells were enlarged at the bottom-left to show the morphology. (**C**) Scratch responses over 30 min with HA (50 µg in 10 µl saline), C48/80 (100 µg in 50 µl saline) and CQ (200 µg in 50 µl saline) injected into nape, respectively. CSF1R depletion significantly decreased the scratch responses to HA (n= 10 for control, n = 11 for CSF1R depletion, p < 0.0001) during the 30 min observation; decreased the responses to C48/80 at both the early (0-3 min, *p* = 0.040) and late (3-30 min, *p* = 0.00011) stages (n = 7 for control, n = 8 for CSF1R depletion); decreased the responses to CQ at both the early (*p* = 0.0049) and late (*p* < 0.0001) stages (n = 10 for control, n = 12 for CSF1R depletion). The secondary test after microglia recovery were done at 14 days after the first one with the same agents. The responses to HA (n = 8, *p* = 0.055), C48/80 (n = 7, *p* = 0.390 for 0-3 min, *p* = 0.125 for 3-30 min) and CQ (n = 11, *p* = 0.938 for 0-3 min, *p* = 0.941 for 3-30 min) were all back to control levels, respectively. \**p* < 0.05, \*\**p* < 0.01, \*\*\**p* < 0.001, \*\*\*\**p* < 0.0001, un-paired t-test. Data were presented as mean ± SEM.

To examine the effect of microglia and macrophage depletion on acute itch transmission, we first tested the itch responses in the HA model and the HA dependent compound 48/80 model in the depletion mice and the control CSF1R^f/f^ mice that received the same doses of TM treatment. In the HA model, the behavioral responses (19.64 ± 2.588 scratch bouts within 30 min) of the depletion mice were significantly less than that of the control mice (51.70 ± 4.069 scratch bouts within 30 min, Fig. 1C). The C48/80 is a very strong itching mediator that act on mast cells and cause degranulation and HA release (McNeil et al., 2015). The scratch responses to C48/80 were much stronger than that to HA treatment in the normal control mice. The scratch frequencies were progressively increased after 3 min and reached the peak within 15 min (Supplementary Fig. 1A). Thus, we separated the total 30 min scratch responses as early stage(0-3in) and late stage (3-30min). As shown in Fig. 1C, in the depletion mice, the scratch bouts were significantly decreased in both the early stage (*p* = 0.040) and the later stage (*p* = 0.00011). The results suggested that the HA dependent itch responses required the CX3CR1+ cells.

We then examined another HA non-dependent model, the CQ model, which induces strong itch responses. CQ treatment in the normal control mice characterized by typical two-phase responses. The mice began scratch immediately after the CQ injection and then calmed down briefly in 3 min. After that, the scratch frequencies increased progressively and reach the peak at around 15 min (supplementary Fig. 1B). In the depletion mice, as shown in Fig. 1C, the scratch bouts were significantly less than the control in both the early (*p* = 0.0049) and late (*p* < 0.0001) stages. The results suggested that the non-histamine dependent itch responses to CQ also required the CX3CR1+ cells.

We also examined the itch responses to β-alanine and mechanical stimulus. β-alanine is a relatively weaker pruritoceptive mediator compared with C48/80 and CQ that the signal is relayed by the NP1 type DRG neurons. Interesting, the scratch responses to β-alanine injection to neck skin in the depletion mice were comparable to that in the control mice (*p* = 0.834) (Fig. S2A). Therefore, β-alanine induced itch sensation transmission did not require microglia or macrophages.

For the mechanical itch test, the mechanical stimuli on the skin behind ear were applied using the Von Frey filaments ranged from 0.02 g to 0.4 g. Similar as previous reported(Pan et al., 2019), the normal control mice were most sensitive to 0.07 g (0.7 N) stimulus. In the depletion mice, the scratch responses to all the tested stimulus strengths were similar as that in the controls, respectively (*p* = 0.544 for group effect with two-way ANOVA, repeated measurement) (Fig. S2B). Therefore, mechanical stimuli induced itch sensation transmission did not require microglia or macrophages. With the above pruritoceptive reagents screening tests, we concluded that the CX3CR1+ microglia and/or macrophages were involved in the itch sensation transmission that elicited by agents depending on HA pathway and the HA non-dependent agent, CQ. The non-affected itch responses to β-alanine and mechanical stimuli suggested that the microglia/macrophage depletion did not affect the behavioral expression of itch responses.

To further confirm whether the transient microglia/macrophage depletion affect the itch transmission pathway in a long term, we did the second tests in the depletion mice for the HA, C48/80 and CQ model at 14 days after the first test (Recovery group in Fig. 1). During the 14 days, no further TM was injected. Thus, microglia and macrophages were repopulated and reached the pre-ablation level (Fig. 1A-B). The results showed that, for all the 3 models, the scratch responses in the mice recovered from the microglia/macrophages depletion were restored and comparable to that in the controls, respectively (Fig. 1C). Therefore, the transient microglia/macrophage depletion would not disrupt itch transmission circuits permanently.

### Depletion of central microglia inhibited acute itch responses to HA, C48/80 and CQ

Since the CSF1R conditional knockout strategy depleted microglia and macrophages at the same time, we could not distinguish the role of central microglia from peripheral macrophages. To dissect the exact role of central microglia in itch transmission, we used the ROSA^iDTR/+^;CX3CR1^CreER/+^ to ablate central microglia alone. The hematopoietic originated monocytes/macrophages have a quick turn-over rate. Several weeks after TM treatment, the circulating monocytes and part of tissue macrophages would be replaced by new cells originated from hematopoietic stem cells (HSCs). Since HSCs do not express CX3CR1, the new cells would not express the induced diphtheria toxin receptors (DTR) and not be affected by the diphtheria toxin treatment(Parkhurst et al., 2013; Peng *et al*., 2016). Thus, we waited 3 weeks after the TM treatment (150 mg/kg, i.p., 4 doses with 48 hr interval), then 2 doses of DT (0.75 μg per mice with 48 hr interval, i.p.) were given. One day after the second DT injection, 76.3±5.3% of Iba1+ cells in the spinal cord were ablated, while 37.9± 7.5% of iba1+ cells in the DRG were ablated as well (Fig. 2B). Thus, the central microglia were efficiently ablated, but the peripheral DRG macrophages were partially ablated as well.

**Figure 2.**
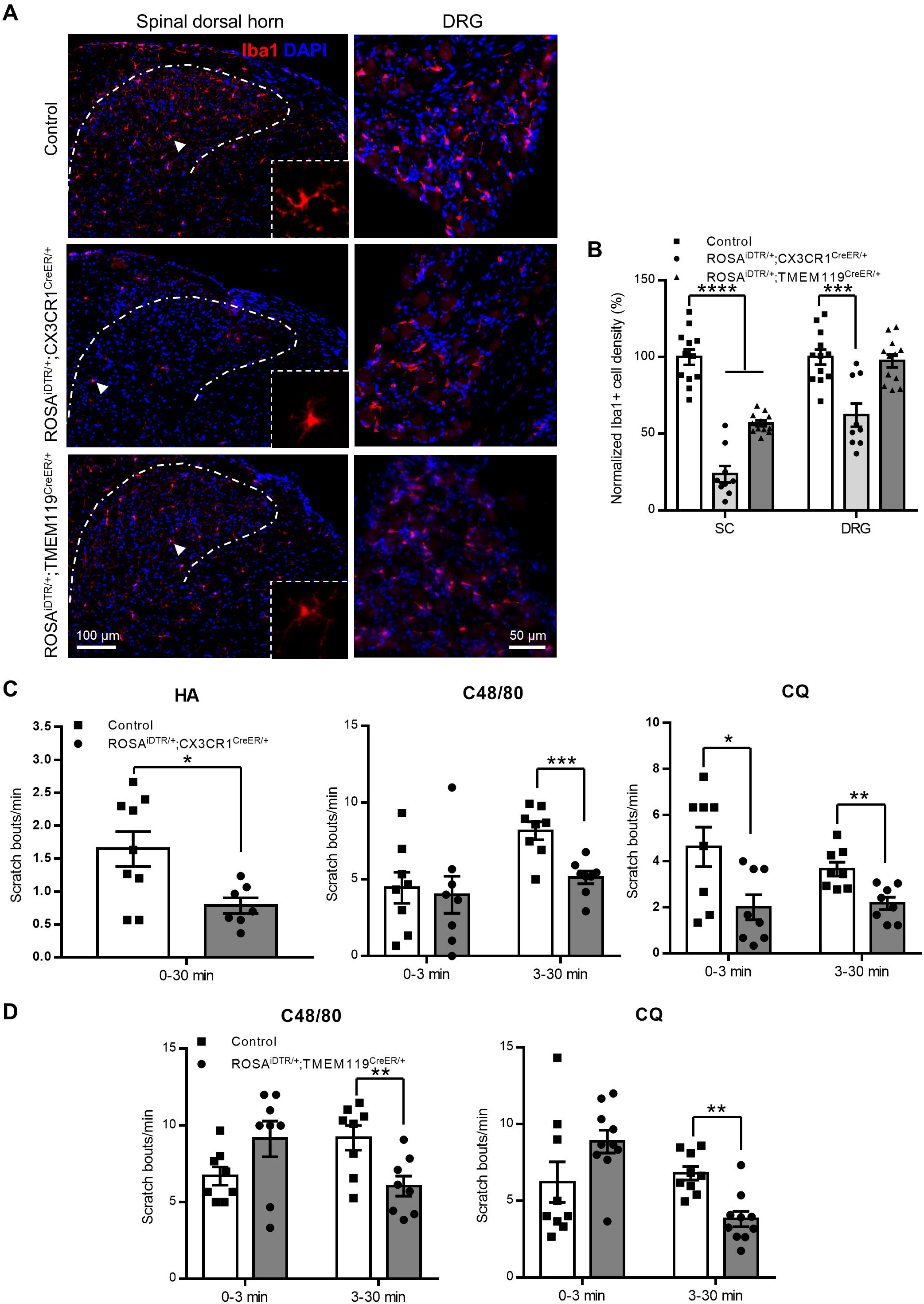
Central microglia ablation inhibited acute itch responses to HA, C48/80 and CQ. (**A-B**) With two doses of Diphtheria Toxin (DT, i.p.,0.75 µg in 200 µl PBS per injection, 48 hr interval) treated to the TM pre-treated ROSA^iDTR/+^;CX3CR1^CreER/+^ and ROSA^iDTR/+^;TMEM119^CreER/+^ mice, microglia cell densities in the SC were reduced to 23.7±5.3% (*p* < 0.0001) and 56.8±1.8% (*p* < 0.0001) of control level, respectively; in the DRG, the Iba1+ macrophages were reduced to 62.1±7.5% in the CX3CR1-CreER mice (*p* = 0.00031), and were not affected in the TMEM119-CreER mice (*p* = 0.709). (n = 3 mice for each group, ROSA^iDTR/+^ mice were used as control and received the same doses of TM and DT, respectively). Arrow indicated cells were enlarged in the bottom right corners. (**C**) Microglia ablation in the ROSA^iDTR/+^;CX3CR1^CreER/+^ mice significantly reduced the scratch responses to HA (n = 9 for control, n = 7 for ablation, p = 0.017) during the 30 min observation; reduced the responses to C48/80 at the late (p = 0.00073), but not early (p = 0.775) stage (n = 8 for control, n = 8 for ablation); reduced the responses to CQ at both the early (*p* = 0.0217) and late (*p* = 0.0026) stages (n = 8 for control, n = 8 for ablation). (**D**) Microglia ablation in the ROSA^iDTR/+^;TMEM119^CreER/+^ mice significantly reduced the scratch responses to C48/80 at the late (p = 0.0091), but not early (p = 0.085) stage (n = 8 for control, n = 8 for ablation); reduced the responses to CQ at the late (p = 0.00038), but not early (p = 0.092) stage (n = 9 for control, n = 10 for ablation). \**p* < 0.05, \*\**p* < 0.01, \*\*\**p* < 0.001, un-paired t-test. Data were presented as mean ± SEM.

To separate the central microglia from the peripheral macrophages/monocytes more clearly. We further used the TMEM119-CreER transgenic mouse to generate the TMEM119^CreER/+^;ROSA^iDTR/+^ mice. Tmem119 gene was found to be specifically expressed in central microglia only(Satoh et al., 2016). We confirmed the tmem119 expression pattern with the TMEM119-EGFP mice. The GFP signals were seen in central microglia cells, but not in DRG macrophages (Fig. S3). With a series of pilot tests, we finally chose a 10-doses TM (150 mg/kg with 48 hr interval, i.p.) injection protocol to achieve a reliable microglia depletion efficiency. 6 days after the last TM injection, two doses of DT (0.75 μg per mice with 48 hr interval, i.p.) were injected to ablate microglia cells. As shown in Fig. 2A-B, Iba1+ cells were reduced by 42.2±1.8% of control in the spinal cord. Most of the remained microglia cells showed ramified morphology similar as the WT control, suggesting that those cells were not affected by DT and the TM induced gene modification was not succeeded in the remained microglia cells. The Iba1+ cells in the DRG were comparable to that of the control mice (*p* = 0.716). Therefore, using the TMEM119-CreER tools, we could specifically manipulate the central microglia without affecting the peripheral macrophages, although the efficiency was less than that of the CX3CR1-CreER.

To examine how the acute itch transmission was affected with central microglia depletion. We tested the acute itch responses in both the DT treated ROSA^iDTR/+^;CX3CR1^CreER/+^ and ROSA^iDTR/+^;TMEM119^CreER/+^ mice (Fig. 2C-D). With HA treatment, the total scratch bouts within 30 min in the ROSA^iDTR/+^;CX3CR1^CreER/+^ (*p* = 0.017) mice were significantly less than that of the control group, which received the same doses of TM and DT treatments. With C48/80 treatment, the scratch responses in the early stage (0-3 min) were not altered in both the ROSA^iDTR/+^;CX3CR1^CreER/+^ (*p* = 0.775) and ROSA^iDTR/+^;TMEM119^CreER/+^ (*p* = 0.085) mice comparing to each control group, but the scratch responses in the late stage (3-30 min) were significantly decreased in both the ROSA^iDTR/+^;CX3CR1^CreER/+^ (*p* = 0.0007) and ROSA^iDTR/+^;TMEM119^CreER/+^ (*p* = 0.0091) mice comparing to each control group. With CQ treatment, the scratch responses in the early stage (0-3 min) were significantly decreased in the ROSA^iDTR/+^;CX3CR1^CreER/+^ (*p* = 0.0217) but not ROSA^iDTR/+^;TMEM119^CreER/+^ (*p* = 0.092) mice comparing to each control group, and the scratch responses in the late stage (3-30 min) were significantly decreased in both the ROSA^iDTR/+^;CX3CR1^CreER/+^ (*p* = 0.0026) and ROSA^iDTR/+^;P2Y12^CreER/CreER^ (*p* = 0.0004) mice comparing to each control group. The results suggested that the central microglia were involved in the HA dependent and non-dependent (CQ) acute itch transmission. The different impact on the early-stage responses among the CSF1R depletion strategy and the two kinds of iDTR strategy suggested that the peripheral macrophages could also contribute to both the HA dependent and non-dependent acute itch transmission, at least in the early stage.

### Microglia were involved in the primary HA, but not CQ itch signal transmission from DRG to spinal cord

To dissect how microglia regulated the neuronal circuits for itch in the spinal level, we examined the c-fos mRNA expression changes at 90 min after the itch agent application via RNAscope. The spinal Npr1+ inter-neurons were known to directly receive both HA dependent and non-dependent itch signals from DRG primary projection (Mishra and Hoon, 2013). And the spinal Sst+ inter-neurons that relay inhibiting signal to DYN+ neurons and work as a disinhibiting function for the itch signals, also occupy an up-stream position in the spinal level of itch transmission (Chen and Sun, 2020; Fatima et al., 2019). Therefore, we co-labeled these neurons together with c-fos. As expected, with HA, C48/80 and CQ treatment, the c-fos positive cell numbers were significantly increased in the cervical spinal dorsal horn of all three WT groups (*p* < 0.0001 for all three groups comparing to naïve) (Fig. 3-5). In the ROSA^iDTR^;CX3CR1^CreER/+^ mice with microglia ablation, HA and C48/80 induced c-fos up-regulation were significantly inhibited compared with the WT groups, respectively (*p* < 0.0001 for both) (Fig. 3B, 4B). However, CQ induced c-fos up-regulation were not altered by microglia ablation (*p* = 0.073) (Fig. 5B). For the Npr1+ neurons, the percentage of c-fos+ cells were increased from 8.60±0.66% to 23.50± 2.05% (*p* < 0.0001) in the HA treated WT mice (Fig. 3C), to 19.85±1.97% (*p* < 0.0001) in the C48/80 treated WT mice (Fig. 4C), to 26.7±4.09% (*p* < 0.0001) in the CQ treated WT mice (Fig. 5C). Microglia ablation significantly decreased HA (*p* = 0.00015, Fig. 3C) and C48/80 (*p* = 0.0021, Fig. 4C), but not CQ (*p* = 0.7460, Fig. 5C) induced c-fos expression in the Npr1+ neurons. For Sst+ neurons, the percentage of c-fos+ cells were increased from 6.36±0.64% to 13.10±1.21% (*p* < 0.0001) in HA treated WT mice (Fig. 3C), to 8.56±0.68% (*p* = 0.0263) in C48/80 treated WT mice (Fig. 4C), to 16.91±2.38 (*p* < 0.0001) in CQ treated WT mice (Fig. 5C). Microglia ablation significantly decreased HA (*p* = 0.0315, Fig. 3C) and C48/80 (*p* = 0.0004, Fig. 4C), but not CQ (*p* = 0.2961, Fig. 5C) induced c-fos expression in the Sst+ neurons as well. These results suggested that spinal microglia were involved in the HA dependent, but not CQ elicited itch signal primary transmission from DRG to the spinal cord.

**Figure 3.**
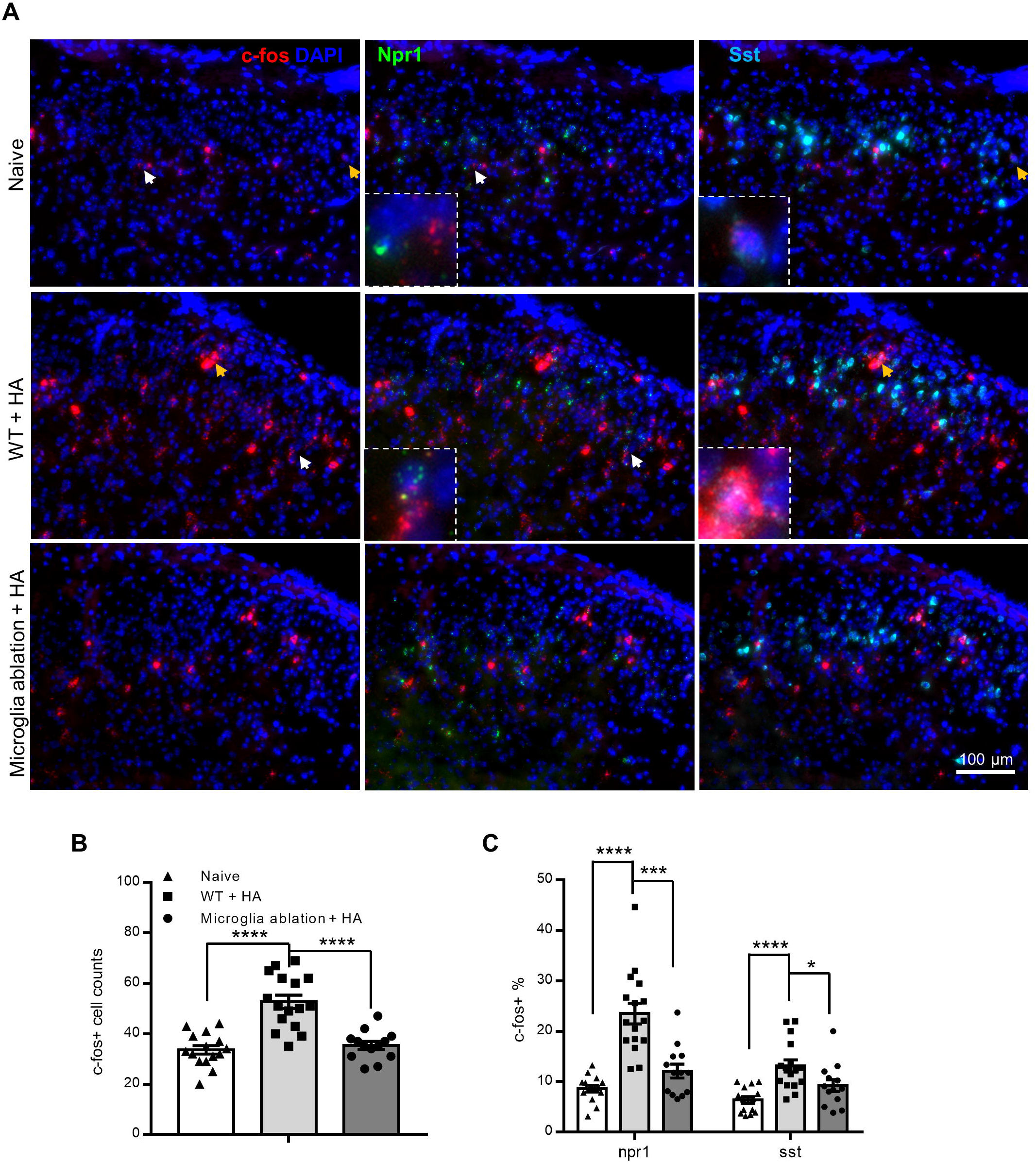
HA induced spinal c-fos mRNA expression was reduced in microglia ablation mice. (**A**) representative RNAscope images showed triple labeling of c-fos, npr-1 and Sst mRNA in the spinal dorsal horn of WT naïve, HA treated WT and microglia ablation (ROSA^iDTR^;CX3CR1^CreER/+^) mice. White and orange arrows indicated c-fos+ cells were enlarged at the bottom-left of the middle and right panels to show the co-labeling with npr-1 and Sst, respectively. (**B-C**) Statistic data showed that HA induced increase of overall c-fos+ cell number in WT mice were greatly reduced in the microglia ablation mice (**B**, *p* < 0.0001 for both naïve vs. WT + HA and WT + HA vs. ablation + HA), and the increase of c-fos+ percentage of Npr1+ and Sst+ cells were also significantly reduced (**C**, *p* < 0.0001 for naïve vs. WT + HA in both Npr1+ and Sst+ cells, *p* = 0.00015 for WT + HA vs. ablation + HA in Npr1+ cells, *p* = 0.0315 for WT + HA vs. ablation + HA in Sst+ cells). n = 15, 16 and 13 images for naïve, WT + HA, and ablation + HA group, respectively. Samples were obtained from 3 mice for each group. \**p* < 0.05, \*\*\**p* < 0.001, \*\*\*\**p* < 0.0001, un-paired t-test. Data were presented as mean ± SEM.

**Figure 4.**
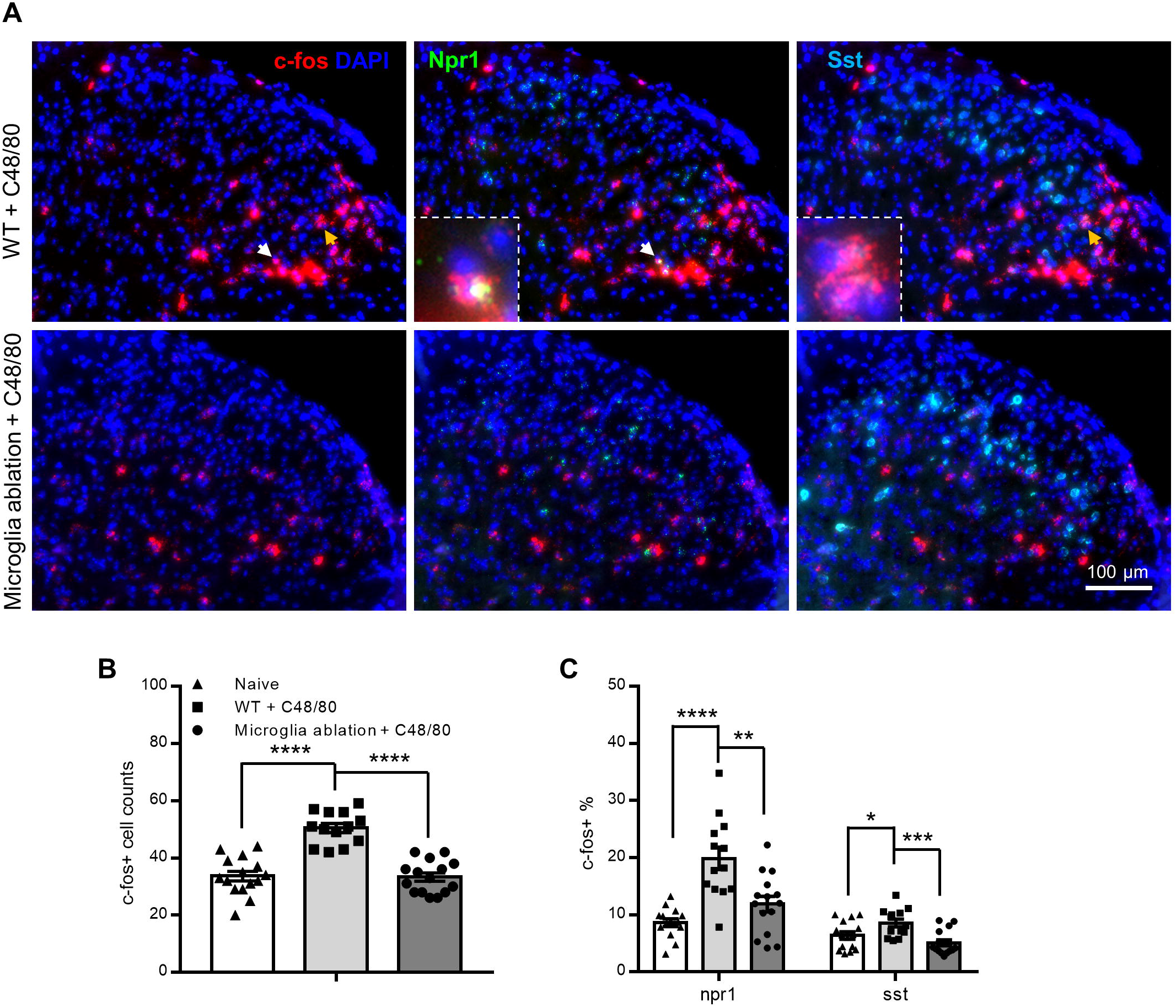
C48/80 induced spinal c-fos mRNA expression was reduced in microglia ablation mice. (**A**) representative RNAscope images showed triple labeling of c-fos, npr-1 and Sst mRNA in the spinal dorsal horn of C48/80 treated WT and microglia ablation (ROSA^iDTR^;CX3CR1^CreER/+^) mice. White and orange arrows indicated c-fos+ cells were enlarged at the bottom-left of the middle and right panels to show the co-labeling with Npr1 and Sst, respectively. (**B-C**) Statistic data showed that C48/80 induced increase of overall c-fos+ cell number in WT mice were greatly reduced in the microglia ablation mice (**B**, *p* < 0.0001 for both naïve vs. WT + C48/80 and WT + C48/80 vs. ablation + C48/80), and increase of c-fos+ percentage of Npr1+ and Sst+ neurons were also significantly reduced (**C**, *p* < 0.0001 for naïve vs. WT + C48/80 in Npr1+ and p = 0.0263 in Sst+ cells, p = 0.00213 for WT + C48/80 vs. ablation + C48/80 in Npr1+ cells, p = 0.00040 for WT + C48/80 vs. ablation + C48/80 in Sst+ cells). n = 13 and 15 images for WT + C48/80, and ablation + C48/80 group, respectively. Naïve data were equal to Fig. 3. Samples were obtained from 3 mice for each group. \**p* < 0.05, \*\**p* < 0.01, \*\*\**p* < 0.001, \*\*\*\**p* < 0.0001, un-paired t-test. Data were presented as mean ± SEM.

**Figure 5.**
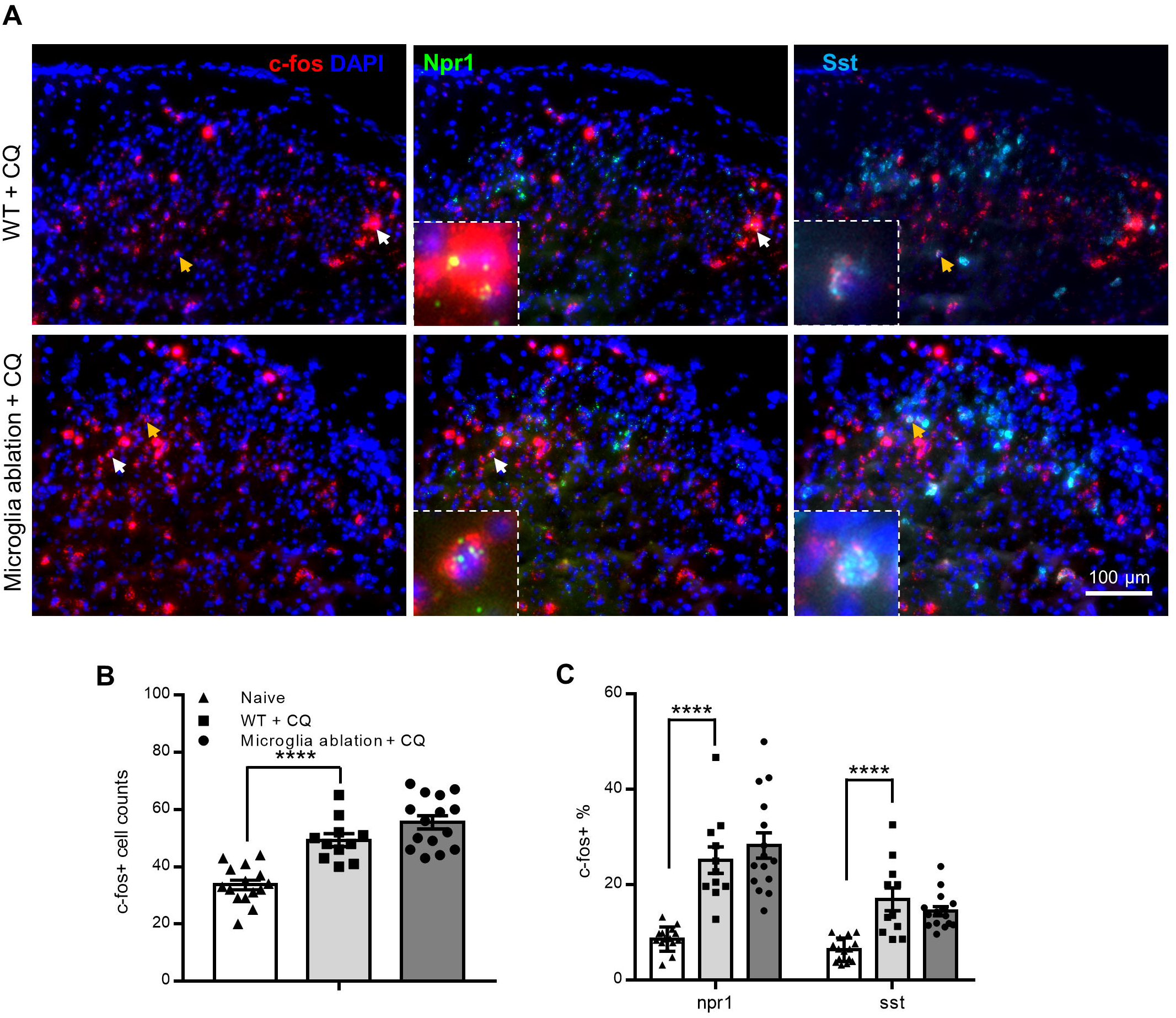
CQ induced spinal c-fos mRNA expression was not affected by microglia ablation. (**A**) representative RNAscope images showed triple labeling of c-fos, npr-1 and Sst mRNA in the spinal dorsal horn of CQ treated WT and microglia ablation (ROSA^iDTR^;CX3CR1^CreER/+^) mice. White and orange arrows indicated c-fos+ cells were enlarged at the bottom-left of the middle and right panels to show the co-labeling with npr-1 and Sst, respectively. (**B-C**) Statistic data showed that CQ induced increase of overall c-fos+ cell number in WT mice were remained in the microglia ablation mice (**B**, p < 0.0001 for naïve vs. WT + CQ, p = 0.0732 for WT + CQ vs. ablation + CQ), and increase of c-fos+ percentage of Npr1+ and Sst+ neurons were also remained (**C**, p < 0.0001 for naïve vs. WT + CQ in both Npr1+ and Sst+ cells, p = 0.439 for WT + CQ vs. ablation + CQ in Npr1+ cells, p = 0.296 for WT + CQ vs. ablation + CQ in Sst+ cells). n = 11 and 15 images for WT + CQ, and ablation + CQ group, respectively. Naïve data were equal to Fig. 3. Samples were obtained from 3 mice for each group. ****p < 0.0001, un-paired t-test. Data were presented as mean ± SEM.

### Spinal CX3CL1-CX3CR1 microglial signal pathway was required for HA dependent, but not CQ elicited itch signal transmission

The above results suggested that spinal microglia were stimulated by very up-stream signals from the HA itch neuronal circuits. Purinergic signal is one of the major components to activate microglia. The P2Y12 receptors are highly expressed in microglia and mediate the microglial process movement towards ATP/ADP gradient (Haynes et al., 2006). Therefore, we first tested the acute itch responses in the P2Y12 KO mice with the HA, C48/80 and CQ models. As shown in Fig. 6, the scratch responses to HA (*p* = 0.891), C48/80 (*p* = 0.572 for early and *p* = 0.184 for late stages) and CQ (*p* = 0.352 for early and *p* = 0.865 for late stages) treatment in the P2Y12 KO mice were all similar as that in the WT control mice, respectively. The results suggested that the P2Y12 receptors were not required for microglia to mediate the acute itch transmission.

**Figure 6.**
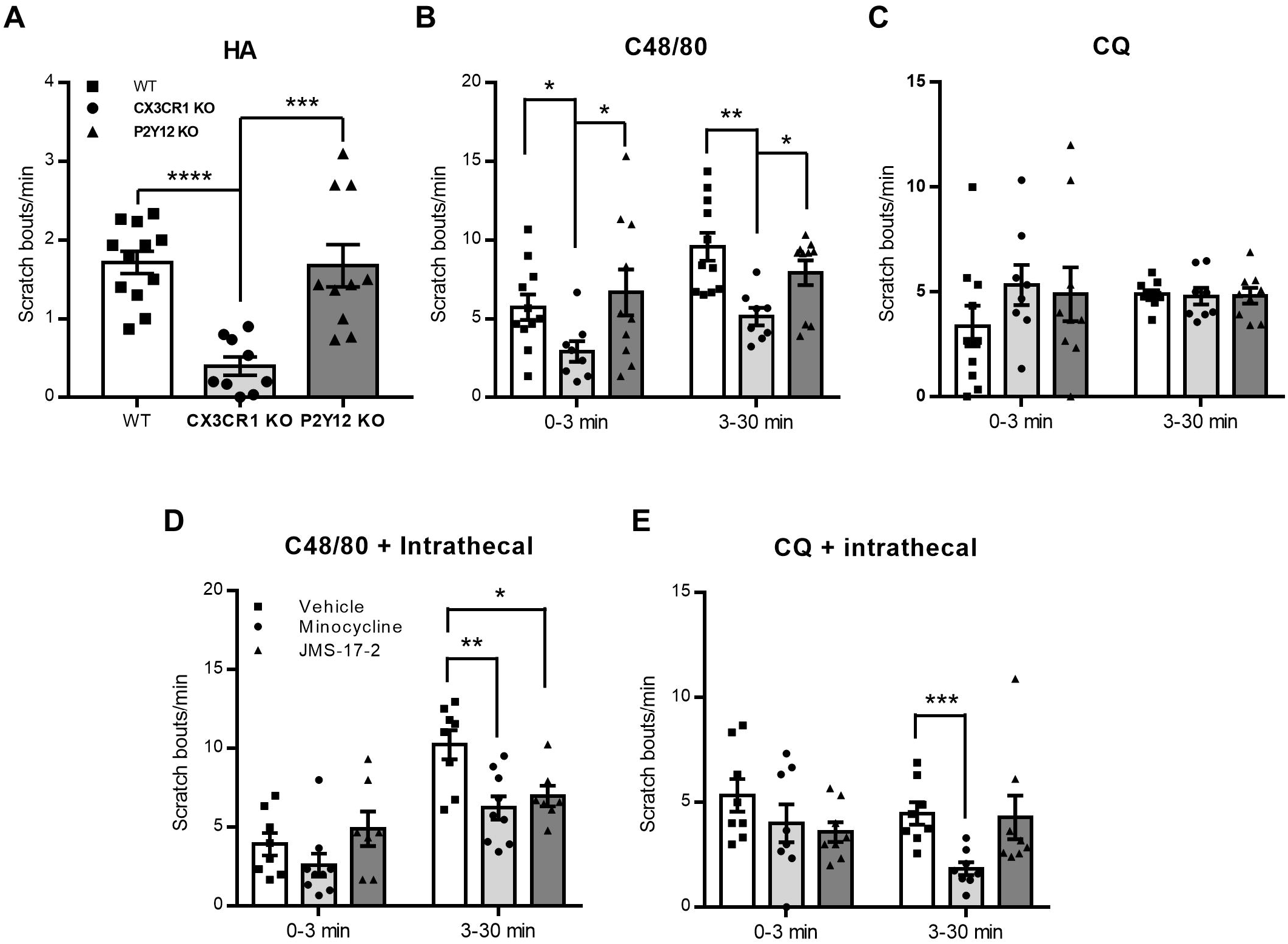
CX3CR1, but not P2Y12 receptor deficit impaired histamine dependent itch responses. (**A**) The Scratch responses to HA in the CX3CR1 KO mice were significantly less than in the WT and P2Y12 KO mice during the 30 min observation (n = 12, 9 and 10 for WT, CX3CR1 KO and P2Y12 KO group, respectively; *p* < 0.0001 for WT vs. CX3CR1 KO, *p* = 0.891 for WT vs. P2Y12 KO). (**B**) The Scratch responses to C48/80 in the CX3CR1 KO mice were significantly less than in the WT and P2Y12 KO mice at both the early (*p* = 0.0206 for WT vs. CX3CR1 KO, *p* = 0.572 for WT vs. P2Y12 KO) and late (*p* = 0.0012 for WT vs. CX3CR1 KO, *p* = 0.184 for WT vs. P2Y12 KO) stages (n = 11, 8 and 10 for WT, CX3CR1 KO and P2Y12 KO group, respectively). (**C**) The Scratch responses to CQ in both the CX3CR1 KO and P2Y12 KO mice were similar as in WT mice at both the early (p = 0.174 for WT vs. CX3CR1 KO, *p* = 0.352 for WT vs. P2Y12 KO) and late (*p* = 0.825 for WT vs. CX3CR1 KO, *p* = 0.865 for WT vs. P2Y12 KO) stages (n = 10, 8 and 9 for WT, CX3CR1 KO and P2Y12 KO group, respectively). (**D**) Intrathecal injection of microglia inhibitor, minocycline (50 μg in 5 μl ACSF) or CX3CR1 antagonist, JMS-17-2 (10 μM in ACSF, 5 μl) to WT mice significantly reduced the late-stage scratch responses to C48/80 (n = 8 for vehicle, n = 9 for minocycline, n = 7 for JMS-17-2; p = 0.0039 for minocycline vs. vehicle, p = 0.015 for JMS-17-2 vs. vehicle). (**E**) Intrathecal injection of microglia inhibitor, minocycline (50 μg in 5 μl ACSF), but not CX3CR1 antagonist, JMS-17-2 (10 μM in ACSF, 5 μl) to WT mice significantly reduced the late-stage scratch responses to CQ (n = 8 for vehicle, n = 8 for minocycline, n = 8 for JMS-17-2; *p* = 0.00071 for minocycline vs. vehicle, *p* = 0.873 for JMS-17-2 vs. vehicle). \**p* < 0.05, \*\**p* < 0.01, \*\*\**p* < 0.001, \*\*\*\**p* < 0.0001, un-paired t-test. Data were presented as mean ± SEM.

The fractalkine or CX3CL1-CX3CR1 signal pathway is another well-known one to mediate the microglial-neuronal interaction and is involved in neuropathic pain and synaptic plasticity (Paolicelli et al., 2011; Zhuang et al., 2007). We then tested the acute itch responses in the CX3CR1 KO mice with the HA, C48/80 and CQ models. The results in Fig. 6 showed that the scratch responses to HA (*p* < 0.0001) in the CXCR1 KO mice were significantly reduced compared to WT control, the responses to C48/80 were also significantly reduced in both early (*p* = 0.0206) and late (*p* = 0.0012) stages. However, the responses to CQ (p = 0.1737 for the early and *p* = 0.8253 for the late stages) were not altered and were comparable to the WT control in both stages. These results suggested that the CX3CL1-CX3CR1 signal pathway was required for microglial activation to mediate the HA dependent itch transmission, but this pathway is not necessary for CQ induced itch responses.

To confirm that the spinal level of microglia anticipated in the acute itch transmission, we examined the effects of intrathecal administration of microglia inhibitor, minocycline, and CX3CR1 antagonist, JMS-17-2 on acute itch transmissions. With minocycline (50 μg in 5 μl ACSF, i.t.) treatment at 30 min prior to the itch agent applications, the scratch responses to both C48/80 and CQ were significantly reduced in the late stage (*p* = 0.0039 for C48/80, *p* = 0.0007 for CQ), but not in the early stage (*p* = 0.219 for C48/80, *p* = 0.282 for CQ) comparing with vehicle controls. With JMS-17-2 (10 μM in ACSF, 5 μl, i.t.) treatment at 30 min prior to the itch agent applications, the scratch responses to C48/80 were significantly reduced in the late stage (*p* = 0.0152), but not early stage (*p* = 0.453); the scratch responses to CQ were not altered in any stage (*p* = 0.076 for the early stage, *p* = 0.873 for the late stage) comparing with vehicle controls (Fig. 6D-E). The results confirmed that spinal microglia were involved in the late-stage itch transmission of both the HA dependent and non-dependent (CQ) types, and spinal microglial CX3CR1 pathway was recruited to the HA dependent itch transmission, but not CQ induced itch transmission.

To further study how the CX3CL1-CX3CR1 signal pathway affect the HA itch neuronal circuits. We examined the spinal c-fos activation with RNAscope again in the CX3CR1 KO mice that treated with HA, C48/80 and CQ, respectively, and co-labeled the Npr1 and Sst neurons. As shown in Fig. 7A-F and Supplementary Fig. 4, the total c-fos+ cells in the spinal dorsal horn of HA (*p* < 0.0001) or C48/80 (*p* < 0.0001) treated CX3CR1 KO mice were significantly decreased compared with the WT control mice that received HA and C48/80 treatment, respectively. Consistent with the behavior data, the CQ (*p* = 0.1367) induced c-fos expression in CX3CR1 KO mice was comparable to the WT control that received CQ treatment. For the Npr1+ neurons, the percentage of c-fos+ cells in HA (*p* = 0.00014) and C48/80 (*p* = 0.0143) treated CX3CR1 KO mice were both significantly less than that of the WT control models, respectively; while the percentage of c-fos+ cells in CQ treated CX3CR1 KO mice were comparable to WT control model. For the Sst+ neurons, the percentage of c-fos+ cells in HA (*p* = 0.0049) treated CX3CR1 KO mice were significantly less than that of the WT control model, but C48/80 (*p* = 0.4188) induced c-fos+/Sst+ neurons were not significantly reduced. The CQ (*p* = 0.1552) induced c-fos+/Sst+ neurons were not affected by CX3CR1 KO. These results suggested that the CX3CL1-CX3CR1 microglial signal pathway played a critical role to promote the primary neuronal responses to HA dependent itch signals in the spinal level, particularly for the Npr1+ neurons.

**Figure 7.**
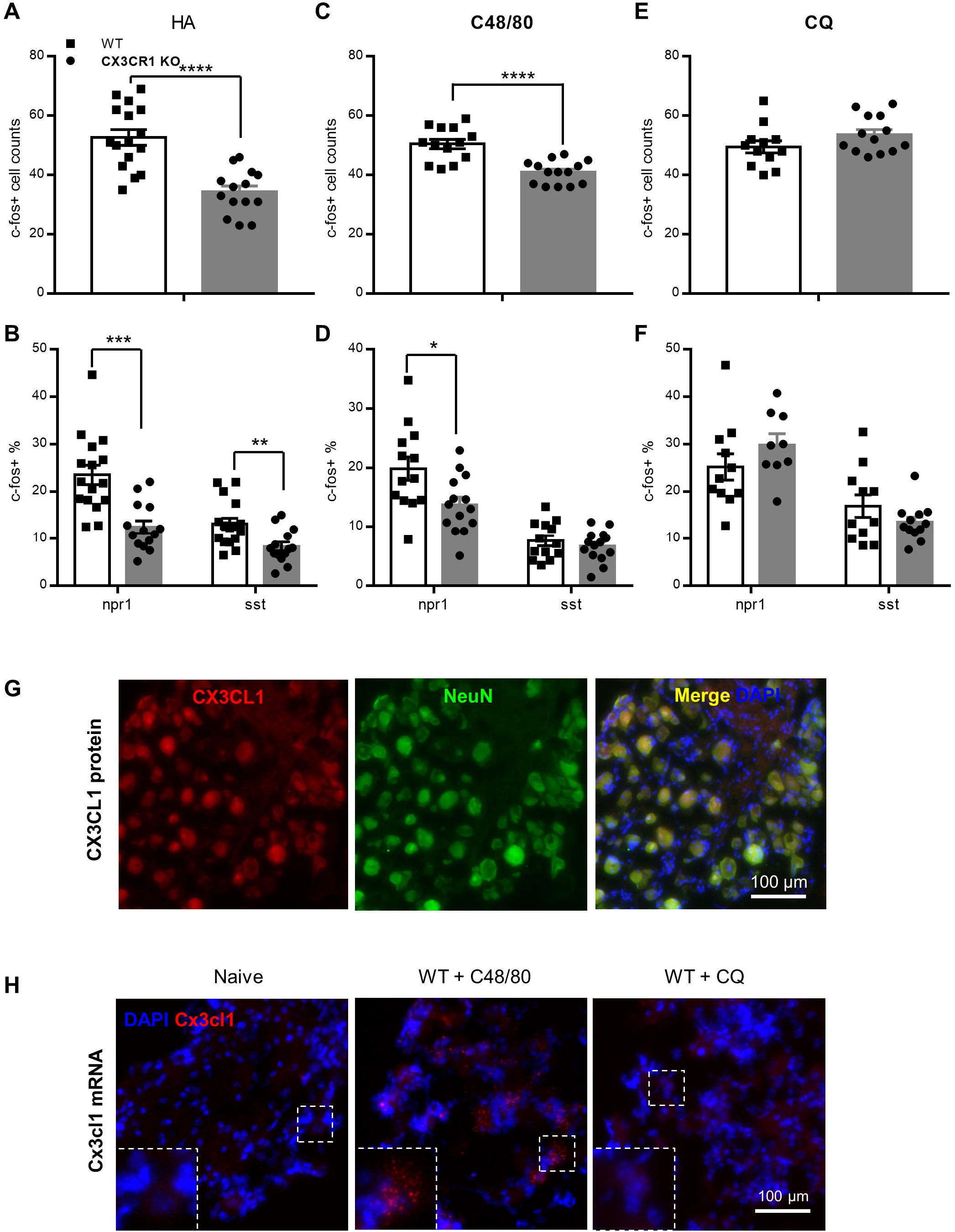
CX3CL1 signal from DRG promoted the primary histamine dependent itch signal transmission via CX3CR1 receptors. (**A-B**) HA induced increase of overall c-fos+ cell number in WT mice were greatly reduced in the CX3CR1 KO mice (p < 0.0001), and the increase of c-fos+ percentage of Npr1+ (p = 0.00014) and Sst+ (p = 0.0049) cells were also significantly reduced (n = 16 for WT, n = 14 for CX3CR1 KO). (**C-D**) C48/80 induced increase of overall c-fos+ cell number in WT mice were greatly reduced in the CX3CR1 KO mice (p < 0.0001), and the increase of c-fos+ percentage of Npr1+ (p = 0.0143), but not Sst+ (p = 0.419) cells were also significantly reduced (n = 13 for WT, n = 14 for CX3CR1 KO). (**E-F**) CQ induced increase of overall c-fos+ cell number in WT mice were remained in the CX3CR1 KO mice (p = 0.137), and the increase of c-fos+ percentage of Npr1+ (p = 0.117) and Sst+ (p = 0.155) cells were also remained (n = 11 for WT, n = 13 for CX3CR1 KO). (**G**) Representative immunostaining images showed CX3CL1 protein expression in DRG neurons of naïve WT mice. (**H**) Representative RNAscope images showed Cx3cl1 mRNA expression in the DRG. Cx3cl1 mRNA was not detectable in the DRG of naïve (n = 10 image samples) WT mice nor CQ (n = 14 image samples) treated (90 min post CQ) WT mice, but was seen in the DRG of C48/80 treated (90 min post C48/80) WT mice (n = 11 of 14 image samples). Samples were obtained from 3 mice for each group. \**p* < 0.05, \*\**p* < 0.01, \*\*\**p* < 0.001, \*\*\*\**p* < 0.0001, un-paired t-test. Data were presented as mean ± SEM.

To investigate where did the CX3CL1 signal come from, we examined the CX3CL1 protein and Cx3cl1 mRNA expression in spinal cord and DRG. Fluorescent immunostaining study revealed that the CX3CL1 protein was constantly presented in the DRG (Fig. 7G) and spinal dorsal horn (Fig. S6A) neuronal cell bodies, and was seen in the spinal nerve fibers in naïve WT mice. the Cx3cl1 mRNA was also constantly expressed in the spinal dorsal horn in naïve WT mice, but was hard to detect any change at 90 min after the itch agent applications (Fig. S6B). However, the Cx3cl1 mRNA was almost non-detectable in naïve DRG. After CQ treatment (90 min), the mRNA signal was still non-detectable in the DRG. After C48/80 treatment (90 min), the Cx3cl1 mRNA expression was clearly seen in DRG neurons (the neuronal locations were recognized by the background morphology). The results suggested that the HA activated DRG sensory neurons could be one of the sources of CX3CL1, and CX3CL1 could be released at the terminal projecting to spinal dorsal horn, although the potential release of CX3CL1 from local neurons in the spinal dorsal horn could not be excluded.

## Discussion

### The neuronal circuit differences for HA and CQ elicited itch signal transmission

The HA and CQ triggered itch signal processing pathways share some common parts in both peripheral DRG and central spinal cord. The CQ receptor, MrgprA3 expressing DRG neurons also express HA receptors and are required for HA itch transmission (Han et al., 2013); NPRA and SST expressing interneurons in spinal cord are required for both HA and CQ signal processing (Fatima *et al*., 2019). However, the pathway differences are critical to define these two types of chemical itch. HA acts directly on H1 and H4 receptors and requires the activation of trpv1 channels (Imamachi et al., 2009). However, DRG neuronal excitation triggered by CQ requires TRPA1, instead of TRPV1 (Wilson et al., 2011). The different activation mechanisms are likely to trigger different neural transmitters release to the spinal cord. For example, glutamate transmission from MrgprA3+ DRG neurons is required for both CQ and HA elicited itch, but NMB from MrgprA3+ DRG neurons is only required for CQ elicited itch (Cui et al., 2022). Therefore, it is possible that there are some un-discovered neural transmitters responsible for certain kinds of itch signal transmission. Here we showed that the CX3CL1 signal was required for HA-dependent itch signal transmission. It is quite possible that CX3CL1 was used as one of the neural transmitters by the DRG neurons to trigger the HA dependent itch transmission.

### The CX3CL1 release mechanism triggered by HA stimulus

The CX3CL1-CX3CR1 pathway was previously reported to be involved in microglial promotion of neuropathic pain (Zhuang *et al*., 2007). CX3CL1 is expressed in both DRG and spinal cord. Peripheral nerve injury was thought to cause the CX3CL1 cleavage and release from DRG neurons (Verge et al., 2004; Zhuang *et al*., 2007). Here we observed an obvious upregulation of Cx3cl1 mRNA in the DRGs with C48/80 stimulus, which triggered endogenous HA release. On the contrary, CQ stimulus did not change the Cx3cl1 mRNA expression. The difference in DRG Cx3cl1 upregulation was correlated with the involvement of CX3CR1 receptors for the itch behavioral responses. Taken together, our results suggested that the HA dependent and non-dependent itch pathways had distinguished downstream responses even at DRG level.

Where was the fractalkine released is still a question remained to be resolved. Central microglia were most likely the major signal receiver of fractalkine, because the reduction of c-fos in spinal Npr1 and Sst neurons and the inhibition of itch responses to HA or C48/80 were seen in both the microglia ablation and CX3CR1 KO mice. However, the peripheral macrophages could also contribute to the enhancement of HA dependent itch signal transmission by responding to fractalkine. In the CSF1R strategy ablation mice or the CX3CR1 KO mice, we observed the inhibition of behavioral response to C48/80 in very early stage (0-3 min), but this phenomenon was not seen in the two kinds of iDTR strategy ablation mice, in which the peripheral macrophages were partially or fully remained. The DRG macrophages are surrounding the neuron bodies and could respond quickly to the fractalkine signal that released from the DRG neuron bodies. The signal added on central microglia could come from the DRG release diffusing to CSF or directly released from DRG projecting terminals. On the other hand, since Cx3cl1 was highly expressed in spinal dorsal horn, the spinal neuronal activities could also trigger fractalkine release.

### Microglial pathways for chemical itch other than CX3CL1-CX3CR1

CQ elicited itch response did not require the CX3CL1-CX3CR1 pathway, but microglia depletion still strongly inhibited the behavioral responses. Although the percentage of c-fos mRNA upregulated Npr1 and Sst neurons were not decreased in the CQ treated microglia depletion mice, we did see an overall c-Fos protein level decrease in spinal dorsal horn with immunostaining (supplementary Fig. 5). The different results between mRNA and protein assays could be due to the different technique sensitivities of RNAscope and immunostaining that RNAscope was more sensitive and would detect and amplify weak c-fos signals. These results suggested that microglia also contributed to the CQ elicited itch signal transmission at spinal level and there were some other signals that triggered microglial responses. For neuropathic pain, the purinergic signal was thought to be important to activate microglia and promote the development of chronic pain. Depletion the purinergic receptors, such as P2X4 (Tsuda et al., 2009), P2X7 (Chessell et al., 2005) or P2Y12 (Tozaki-Saitoh et al., 2008), all dramatically alleviated or totally blocked neuropathic pain. P2X4 can respond to ATP at nanomolar level, which is about one thousand times lower than that for P2X7. Therefore, P2X4 could be a good candidate to mediate the quick microglial responses for itch transmission. P2Y12 receptors are highly expressed in central microglia and mediate the microglial process movement toward ATP gradient (Haynes *et al*., 2006). Unexpectedly, our results showed that the P2Y12 KO mice did not show any deficits in chemical itch response, suggesting that this directional process movement was not necessary for microglia to enhance neuronal excitability quickly. Further studies are required to examine other signal pathways for microglia to promote chemical itch transmission.

The down-stream of microglial responses to itch signals is another question remained to be resolved in the future. Cytokines such as IL-31 and TNF-α have been found to contribute to acute and chronic itch (Cevikbas et al., 2014; Miao et al., 2018). Microglia was one of the major sources of TNF-α and are involved in acute inflammatory pain by enhancing neuronal excitability (Berta et al., 2014). Thus, microglia released TNF-α is a potential mediator for acute chemical itch as well. BDNF is known to be the down-stream of P2X4 that released by microglia to mediate neuropathic pain through disinhibiting mechanism (Beggs et al., 2012). Whether this pathway also contribute to itch signal transmission is worthy to test as well.

In conclusion, our present study revealed that central microglia played a critical role in promoting acute chemical itch signal transmission that induced by HA dependent or non-dependent (CQ) agents. However, microglia participated in the HA dependent and non-dependent itch signal transmission in different ways. For the HA dependent signals, the CX3CL1-CX3CR1 signal pathway could be the major component to trigger microglial responses and then promote the neuronal activities of the spinal Npr1+ and Sst+ neurons. The CX3CL1 signal was most likely to be released by the HA activated DRG sensory neurons that projected to spinal dorsal horn. However, how the CQ signal activate microglia and the down-stream microglial response mechanisms remained unclear.

## Acknowledgements

This work was supported by the Jiangxi Province “2000 talent plan” to JP (jxsq2018106039), the National Natural Science Foundation of China (32060199 to J.P., 82071245 to H.P., and 31971035, 31771182 to B.M.L.) and Jiangxi Province Natural Science Foundation (20171ACB20002 to B.M.L.).

## Author contributions

J.P., H.P. and Y.Y. conceived of the study. Y.Y., B.M., H.X.Z., X.Y., M.T.X. and Y.L. performed experiments and analyzed data. J.P., Y.Y, Y. U. L, H.P., C.L.M. and B.M.L. wrote the manuscript.

## Materials and methods

### Animals

All experimental procedures were approved by the Institutional Animal Care and Use Committee of Nanchang University. We followed the guidelines set forth by the Guide of the Care and Use of Laboratory Animals 8th Edition. The P2Y12 KO mice were originally obtained from Dr. Long-Jun Wu lab at Mayo Clinic, which was originally generated by Dr. Pamela B. Conley(Andre et al., 2003). CX3CR1-CreER (#021160), TMEM119-CreER (#031820), TMEM119-EGFP (#031823), CSF1R-flox (#021212), and ROS26-iDTR (007900) mice were originally purchased from Jackson Laboratory. CX3CR1-CreER mice were crossed with CSF1R-flox mice to get the CSF1R^flox/flox^;CX3CR1^CreER/+^ mice, and crossed with the ROSA-iDTR mice to get the ROSA^iDTR/+^;CX3CR1^CreER/+^ mice. TMEM119-CreER mice were crossed with the ROSA-iDTR mice to get the ROSA^iDTR/+^;TMEM119^CreER/+^ mice. Wild type (WT) C57BL6/J mice were obtained from SLAC laboratory animal CO. LTD (Changsha, China). Mice were group (4-5 per cage) housed in 12/12 light/dark cycle, 23 ± 1 ºC vivarium environment. Food and water were available ad libitum. Mice (8-14 weeks old) were assigned to experimental groups randomly within a litter. Experimenters were blind to drug treatments and mouse genotypes until all data collection was done. Both male and female mice were used.

### Microglia ablation

#### CSF1R strategy

CSF1R^flox/flox^;CX3CR1^CreER/+^ transgenic mice were used for this purpose. Intraperitoneally (i.p.) injection of tamoxifen (TM, 150 mg kg^-1^ in corn oil, 3 doses with 48-hr intervals) were used to trigger the csf1r gene knockout in CX3CR1+ cells. CSF1R^flox/flox^ mice were used as control and received the same doses of TM. Acute itch models were tested at 24 hr after the last TM treatment.

#### iDTR strategies

ROSA^iDTR/+^;CX3CR1^CreER/+^ and ROSA^iDTR/+^;P2Y12^CreER/CreER^ transgenic mice were used. For the ROSA^iDTR/+^;CX3CR1^CreER/+^ mice, 4 doses of TM (150 mg kg^-1^ in corn oil) were i.p. injected with 48-hr intervals to trigger the DTR expression in CX3CR1+ cells. 3 weeks after the last TM injection, two doses of Diphtheria Toxin (DT) were i.p. injected (0.75 µg per mice) with a 48-hr interval to ablate central microglia, but avoided to ablate most of the circulating monocytes and peripheral macrophages. For the ROSA^iDTR/+^;TMEM119^CreER/+^ mice, 10 doses of TM (150 mg kg^-1^ in corn oil) were i.p. injected with 48-hr intervals to trigger the DTR expression in central microglia. 5 days after the last TM injection, two doses of Diphtheria Toxin (DT) were i.p. injected (0.75 µg per mice) with a 48-hr interval to ablate central microglia.

#### Intrathecal injections

The microglia activation inhibitor, minocycline (10 μg/μl in ACSF that contained 0.1% DMSO and 0.4% PEG300) and the CX3CR1 antagonist, JMS-17-2 (10 μM in ACSF that contained 0.1% DMSO and 0.4% PEG300) were intrathecal (i.t.) injected as previous described. In brief, mice were hand restricted, a 31G needle that attached with 10-μL Hamilton syringe (Hamilton Bonaduz AG) were direct lumbar punched between L5 and L6 vertebrae of the spine with around 15 ° angle, successful insertion was indicated by tail flick. 5 μl of the drug solution or control vehicle was injected into the spinal fluid space in 2 min, and the needle was hold in place for one more minute. The i.t. injections were done 30 min prior to the itch agent application.

### Behavioral Testing

#### Acute mechanical itch

To test the acute mechanical itch, the fur behind the ears was shaved 5 days before testing. Mice were habituated for 30 min in behavioral testing apparatus (IITC, Life Science) for 2 consecutive days. On the testing day, mice were placed in the plastic chambers and allowed at least 30 min for habituation. Mice then received five separate mechanical stimuli for 1 s with 3-5 s intervals at randomly selected sites on the skin behind the ears. Mechanical stimuli were delivered with von Frey filaments (0.02-0.16g, North Coast medical). The scratching response of hind paw toward the poking site was considered as a positive response (Pan *et al*., 2019).

#### Acute chemical itch

To test the acute chemical itch, the fur on the neck was shaved 5 days before testing. Mice were habituated same as for the acute mechanical itch test. On the testing day, mice were placed in the plastic chambers and allowed at least 30 min for habituation. Then the behavior of mice was video recorded for at least 30 min after chemical injection. Compound histamine (50 µg, Sigma #H7250) in 10 µl of sterile saline was injected intradermally into the nape. Compound chloroquine (200 µg, Sigma #C6628), β-alanine (50 mM Sigma #146064), C48/80 (100 µg, Sigma #C2313) in 50 µl of sterile saline was injected intradermally into the nape. Scratching bouts were counted for 30 min after injection.

### Immunofluorescence

Experimental mice were deeply anesthetized by 1% pentobarbital sodium (50 mg/kg, i.p.) and transcardially perfused with 0.9% saline followed by 4% paraformaldehyde solution. The cervical segment of spinal cord (C3-C5) and their connected DRGs were collected and post-fixed with the same 4% PFA for 4–6h at 4 °C, and then gradient dehydrated in 20% and 30% (w/v) sucrose solution sequentially. Sample sections (14 μm in thickness) were prepared on gelatin-coated glass slide with a freezing microtome (Leica CM900, Germany). The sections were blocked with 5% goat serum and 0.3% Triton X-100 (Sigma) in TBS buffer for 45 min, and then incubated overnight at 4 °C with primary antibody for rabbit-anti-Iba1 (1:1000, Abcam, Catalogue ab178846), rat-anti-F4/80 (1:500, Biolegend, Catalogue #123102) and rabbit-anti-c-Fos (1:500, Cell Signaling, Catalogue #2250). After rinse for three times for at least 30 minutes with TBS buffer, the sections were then incubated for 90 min at room temperature with secondary antibodies (1:500, Alexa Fluor 568, Alexa Fluor 488, Life Technologies). After three rinses, slices were incubated with DAPI solution for 5 minutes and followed by washout. Fluorescent images were obtained with a fluorescence microscope (EVOS FL Color, life technologies). Cell counting and fluorescent signal intensity was quantified using Image J software (1.52a, National Institutes of Health, Bethesda, MD).

### RNAscope

The frozen section samples of the same sets for the above immunofluorescence were taken. Samples were washed in PBS for 5 min and air dried. Next about 5-8 drops of RNAscope® hydrogen peroxide were added to coat the samples and incubated at room temperature for 10 min and then washed with distilled water. Then immerge the samples into the RNAscope target repair reagent at 98-100 °C for 5 min, then transferred to distilled water for cooling. The samples were then incubated with 100% ethanol for 3 minutes and drawn a hydrophobic ring around it. Appropriate amount of RNAscope® protease III reagent was dropped to completely cover the sections, and incubation was conducted at 40°C for 30 min. The samples were then process immediately with the provided standard RNAscope assay (Advanced Cell Diagnostics, Inc.). mm-fos, mm-npr1 and mm-sst probes were used for triple labeling at opal 690, 520 and 570 channels. mm-cx3cl1 probe was used for single labeling at opal 690 channel. The samples were finally stained with DAPI. The fluorescent images were obtained with the EVOS microscope and analyzed with ImageJ as well.

### Statistical Analysis

Statistical analysis was performed using GraphPad Prism 7.00 (GraphPad software, Inc). Unpaired Student’s test (t-test) and two-way ANOVA with repeated measurement were applied for group-group comparation. *p* < 0.05 was considered statistically significant.

