## supplemental figure 1-6 for "Microglia are involved in regulating histamine dependent and non-dependent itch transmissions with distinguished signal pathways"

### SUPPLEMENTARY FIGURES AND FIGURE LEGENDS

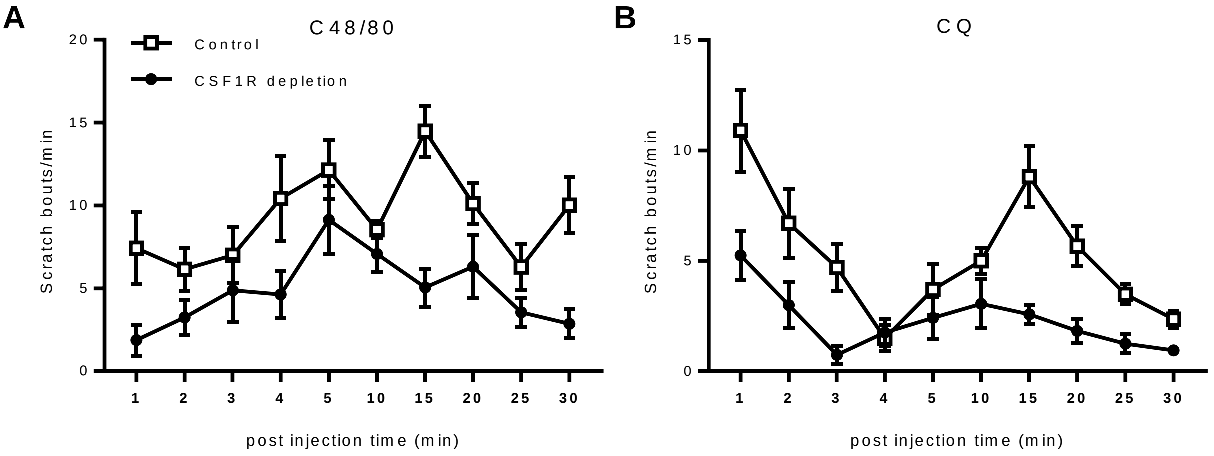


**Figure S1. Timecourse of scratch responses to C48/80 and CQ. (A)** The scratch frequencies were progressively increased and reached the peak within 15 min after C48/80 injection to the control mice. CSF1R depletion significantly decreased the scratch bouts overall ($p$ = 0.0002, F(1, 13) = 26.61 for group effect, two-way ANOVO). **(B)** The scratch responses to CQ showed a typical two-phase pattern that the immediate responses calmed down briefly in 3 min and then grew up and reach the peak at around 15 min after CQ injection to the control mice. CSF1R depletion significantly decreased the immediate responses and flattened the late stage peak ($p$ < 0.0001, F(1, 20) = 25.34 for group effect, two-way ANOVA with repeated measurement).

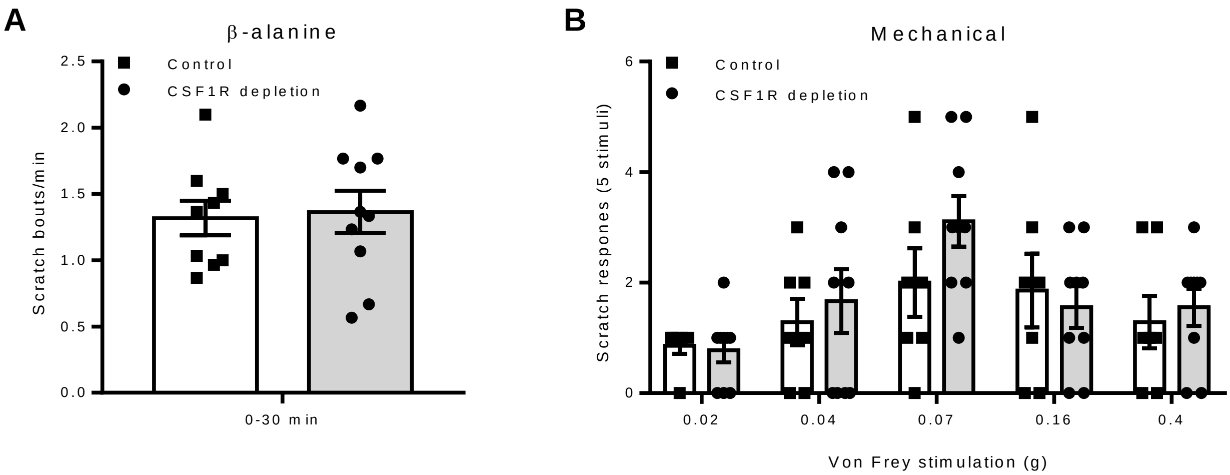


**Figure S2. Microglia depletion did not affect β-alanine induced acute itch and Von Frey induced mechanical itch. (A)** The Scratch responses to β-alanine were not different between CSF1R depletion and control mice during the 30 min observation (n=10 for CSF1R depletion, n=9 for control, $p$ = 0.7343, un-paired t-test). **(B)** the Von Frey induced mechanical itch responses were not affected by CSF1R depletion (n=9 for CSF1R depletion, n=7 for control, $p$ = 0.544, F (1, 14) = 0.387, two-way ANOVA with repeated measurement). Data were presented as mean ± SEM.

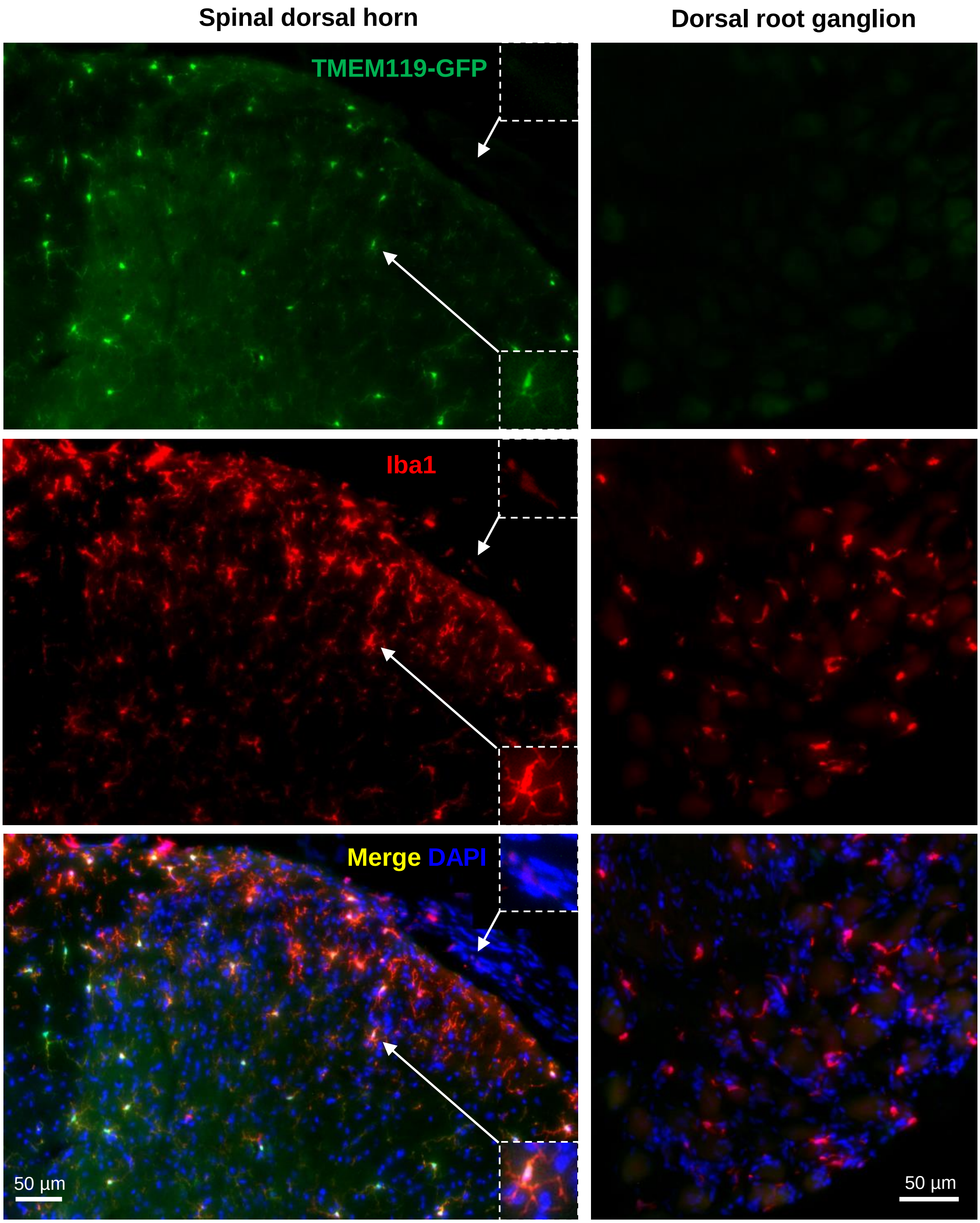


**Figure S3. Specific expression of TMEM119 in central microglia but not peripheral macrophages.** Samples were obtained from TMEM119-EGFP mice. As shown, in the spinal dorsal horn, all GFP labeled cells were Iba1 positive, while some Iba1+ cells in the surface and the attached spinal nerve were GFP negative. In the DRG, all Iba1+ macrophages were GFP negative. The arrow indicated Iba1+/GFP+ microglia cell and Iba1+/GFP- spinal nerve macrophage cell were enlarged in the right bottom and top corners, respectively.

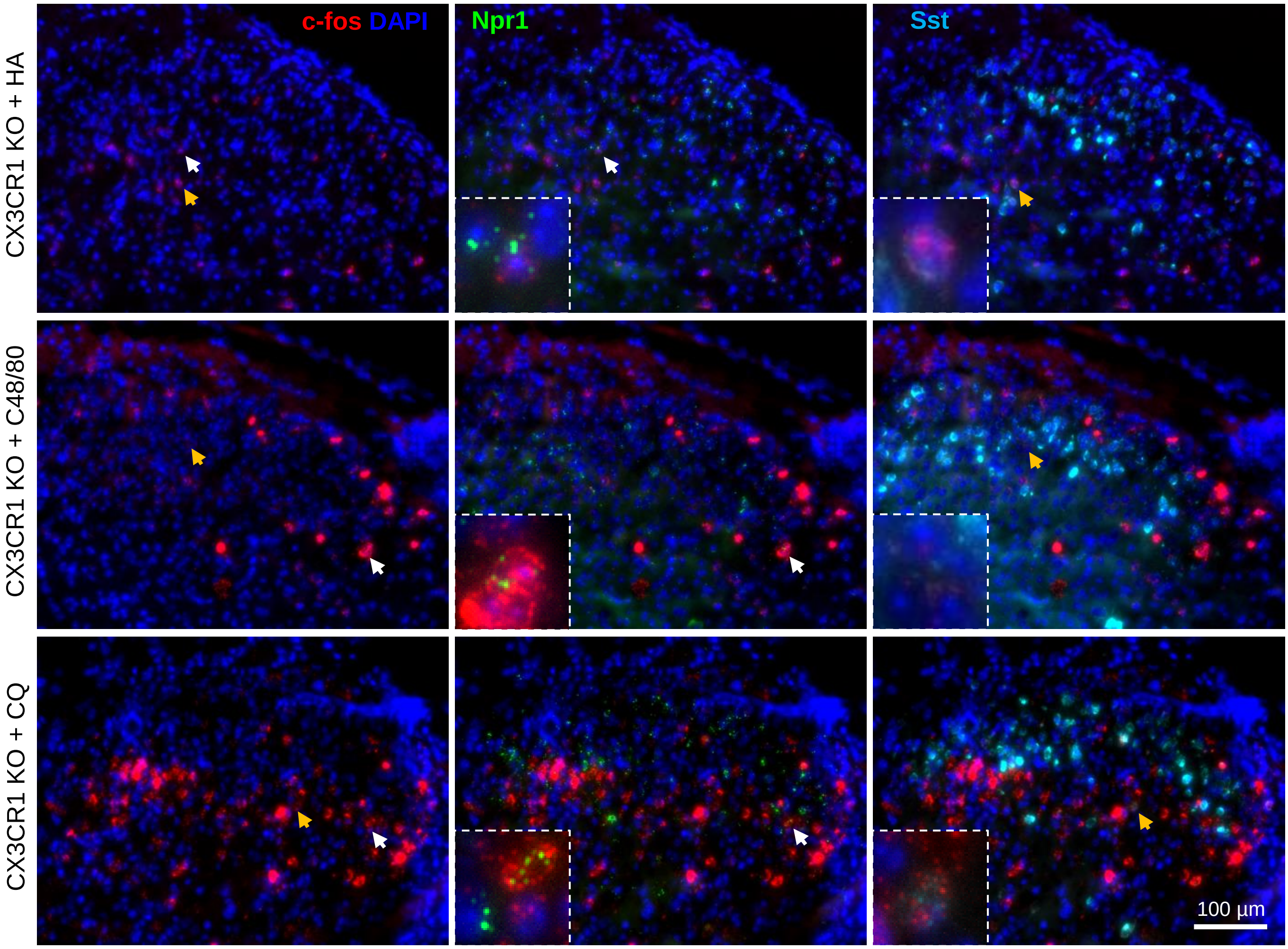


**Figure S4. Representative RNAscope images showed triple labeling of c-fos, npr-1 and sst mRNA in the spinal dorsal horn of CX3CR1 KO mice by HA, C48/80 and CQ treated respectively.** White and orange arrows indicated c-fos+ cells were enlarged at the bottom-left of the middle and right panels to show the co-labeling with npr-1 and sst, respectively.

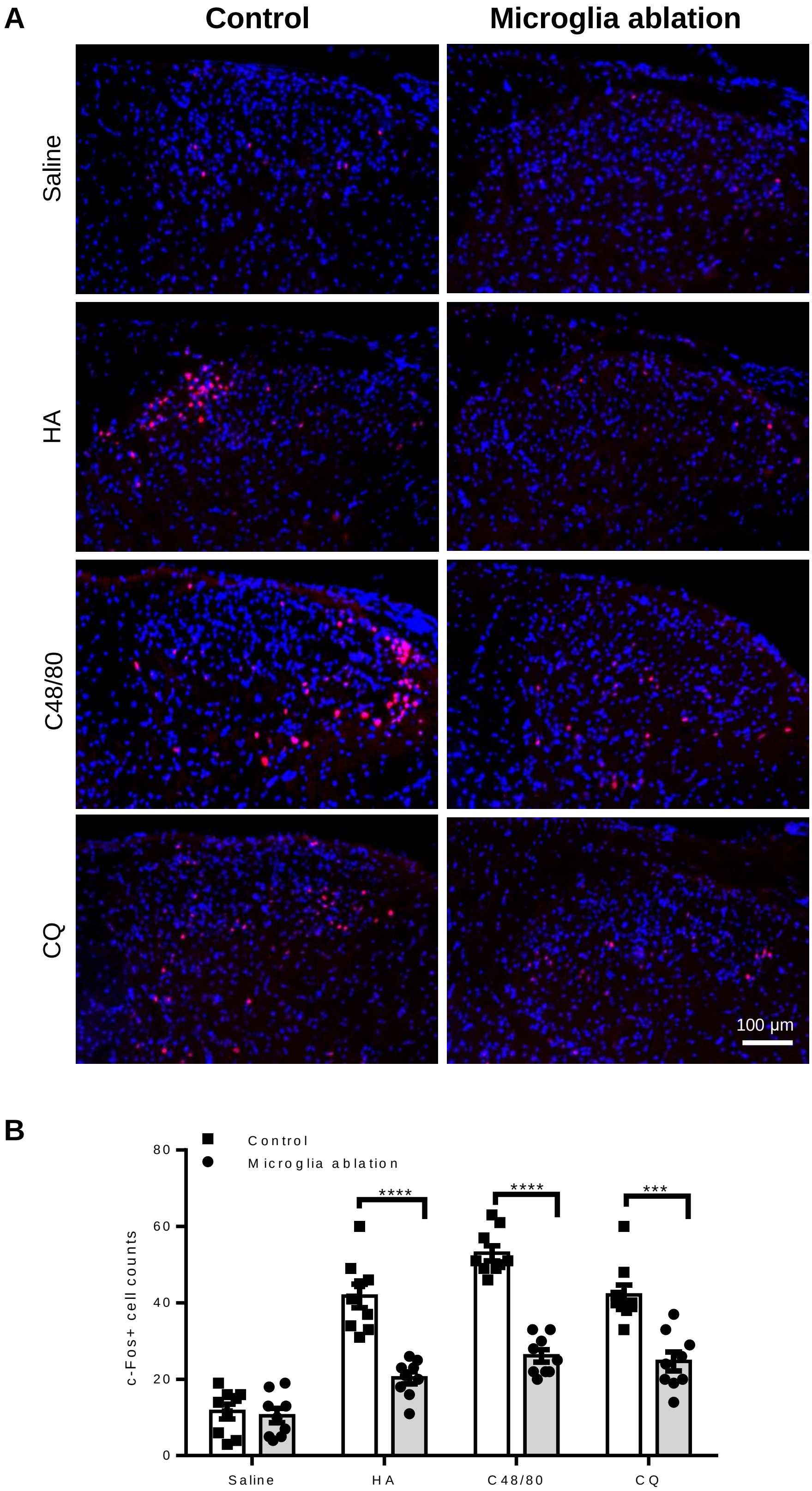


**Figure S5. Central microglia ablation inhibited c-Fos protein expression in the spinal cord induced by HA, C48/80 and CQ. (A)** Immunofluorescence of c-Fos expression in spinal cord of control and microglia ablation (ROSA$^{iDTR/+}$:P2Y12$^{CreER/CreER}$) mice. Samples were obtained at 90 min after the treatment of HA , C48/80 and CQ, respectively. **(B)** Statistic data showed that c-Fos expression was not altered by microglia ablation in the naïve mice ($p$ = 0.688), but was decreased in microglia ablation mice of HA ($p < 0.0001$), C48/80 ($p < 0.0001$) and CQ ($p = 0.00016$) groups. n = 9 images from 3 mice for each group. ***$p < 0.001$, ****$p < 0.0001$, un-paired t-test. Data were presented as mean ± SEM.

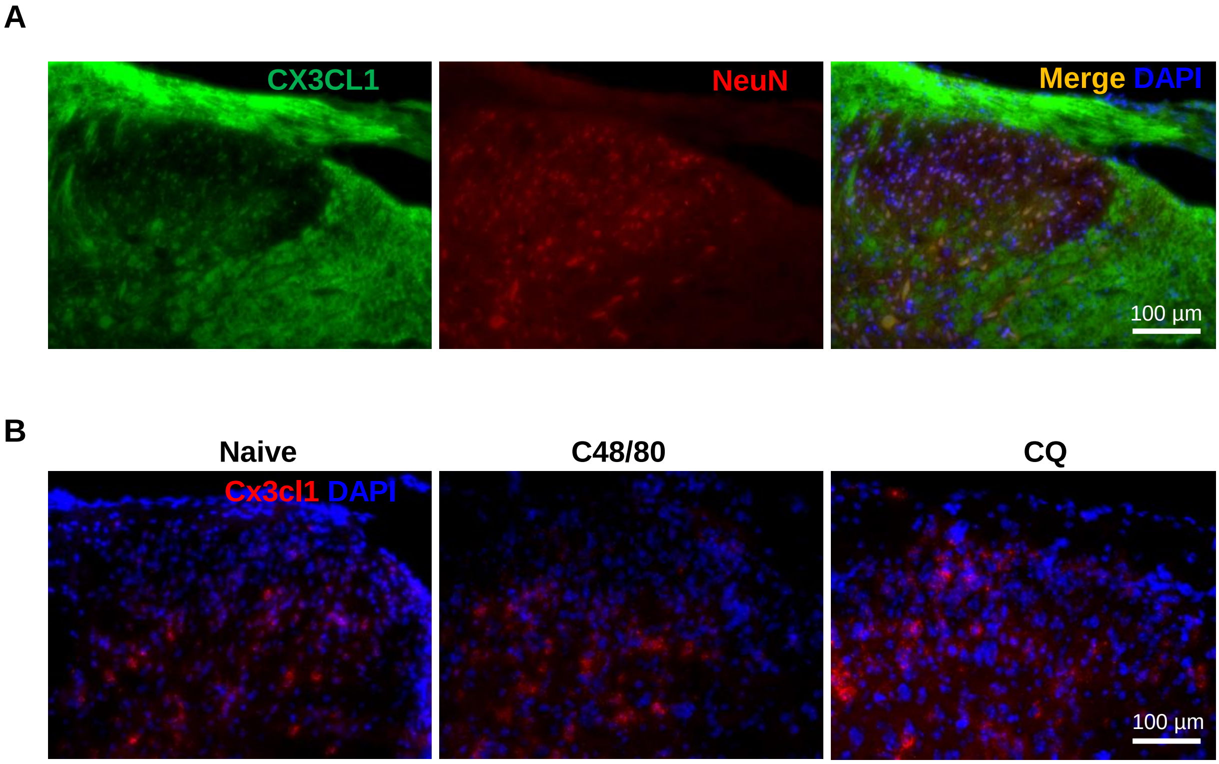


**Figure S6. CX3CL1 was constantly expressed in spinal dorsal horn. A**) Antibody staining showed CX3CL1 protein distribution in the spinal dorsal horn neurons (NeuN+) and spinal nerve fibers. **B**) RNAscope assay showed cx3cl1 mRNA was constantly expressed in in the spinal dorsal horn with or without itch mediator applications.
